# Spatial Transcriptomics Prediction from Histology Images at Single-cell Resolution using RedeHist

**DOI:** 10.1101/2024.06.17.599464

**Authors:** Yunshan Zhong, Jiaxiang Zhang, Xianwen Ren

## Abstract

Spatial transcriptomics (ST) offers substantial promise in elucidating the tissue architecture of biological systems. However, its utility is frequently hindered by constraints such as high costs, time-intensive procedures, and incomplete gene readout. Here we introduce RedeHist, a novel deep learning approach integrating scRNA-seq data to predict ST from histology images at single-cell resolution. Application of RedeHist to both sequencing-based and imaging-based ST data demonstrated its outperformance in high-resolution and accurate prediction, whole-transcriptome gene imputation, and fine-grained cell annotation compared with the state-of-the-art algorithms.

## Introduction

Spatial transcriptomics (ST) is a cutting-edge technology that enables the analysis of gene expression patterns in tissues with spatial resolution. This technology holds tremendous value for biological research, primarily because it bridges the gap between traditional bulk sequencing and single-cell analysis by providing a spatial context to gene expression data. Recently, numerous ST platforms have emerged. Fundamentally, the current ST technologies can be categorized into two primary categories: imaging-based ST and sequencing-based ST. Sequencing-based ST relies on sequencing technology to quantify gene expression at spot level, such as 10x Visium and Slide-seq^1^. This type of approaches enables the capture of whole transcriptomic expression profiles, but it is challenging to achieve single-cell resolution and may result in the omission of crucial information within the spot-spot gap region. In contrast, imaging-based ST, such as MERFISH^2^, Xenium^3^ and CosMx^4^, visualizes and quantifies gene expression directly within intact tissues using microscopy and fluorescent probes. This type of approaches enables the observation of gene expression patterns at single-cell or even subcellular levels, but it is unable to capture whole transcriptomic profiles.

Although ST is a potent tool for investigating the transcriptome, its clinical application is constrained by its high cost and time-consuming nature. In contrast, histology imaging is readily accessible and cost-effective in clinical settings. Hence, establishing a connection between histology images and ST, and ultimately achieving spatial transcriptomic predictions based on histology images, is highly meaningful. Recently, several methods have been developed for this propose, such as Hist2ST^5^, iStar^6^, BLEEP^7^ and ImSpiRE^8^. They employ a transformer-based^9^ deep learning algorithm to extract image features from histology images, and subsequently predict expression based on these features. Despite these approaches can predict transcriptomics, they still have several limitations. First, these methods can only predict the expression of hundreds or thousands of highly variable genes, but they cannot predict the whole transcriptomic expression profiles due to GPU memory limitation. Second, predicting at single-cell resolution is challenging due to the fact that transformers often output patch-level features instead of pixel-level features. Third, current methods often require spatial coordinates of patches as input and establish the association between expression and spatial positions, thereby restricting their applicability in predicting novel images. Fourth, without incorporating single-cell RNA sequencing (scRNA-seq) data, these methods have limitations in accurately predicting spatial transcriptomic expression profiles. Finally, these approaches are designed for analyzing sequencing-based ST, and thus are not suitable for directly analyzing imaging-based ST.

In this study, we introduce RedeHist, a deep learning framework that integrates histological cell segmentation and scRNA-seq data to predict ST at single-cell resolution from histology images, aiming to overcome the existing bottlenecks in the field. First, as many cell segmentation methods have been developed to identify cell or nuclei regions based on histology images, detected transcripts, or both, such as cellpose^10^, VistoSeg^11^, SCS^12^, Baysor^13^ and RedeFISH^14^, the inclusion of cell segmentation in RedeHist not only enhances prediction resolution but also maximizes the utilization of single-cell information, particularly for imaging-based ST data. Second, RedeHist utilizes U-net^15^ to extract pixel-level features and integrates nucleus masks from segmentation results to generate cell-level latent embeddings. Because of the irregular nature of cell morphology, patch-based features, without considering nuclei masked regions, are unsuitable for directly predicting cell expression. Third, inspired by a ST deconvolution method we recently developed, named as Redeconve^16^, we hypothesize that the expression of a spot region in sequencing-based ST could be represented by the summed expression of corresponding single cells. Hence, we implemented a deconvolution-like algorithm using scRNA-seq references to predict cell expression on histology images. Incorporating scRNA-seq references not only improves the accuracy of expression prediction but also enables the imputation of unmeasured genes. In addition, it also streamlines cell annotation by facilitating automatic label transfer. Stringent comparison with the state-of-the-art (SOTA) algorithms including iStar and Hist2ST demonstrates the superiority of RedeHist in terms of prediction accuracy, resolution and gene imputation. RedeHist may serve as a valuable tool for deciphering the spatial organization of gene expression at single-cell resolution based on histology images, with potential implications for understanding complex biological processes and advancing targeted therapies^17,18^.

## Results

### The Architecture of RedeHist

The overall architecture of RedeHist is given in Fig 1. Basically, our approach employs a deep neural network integrated with nuclei segmentation results to predict transcriptomic profiles at single-cell resolution from histology images. RedeHist takes histology images, ST data, and scRNA-seq references as inputs, then generates outputs consisting of single cells identified on the images along with their whole transcriptomic expression profiles, spatial coordinates, and annotations.

**Fig 1.**
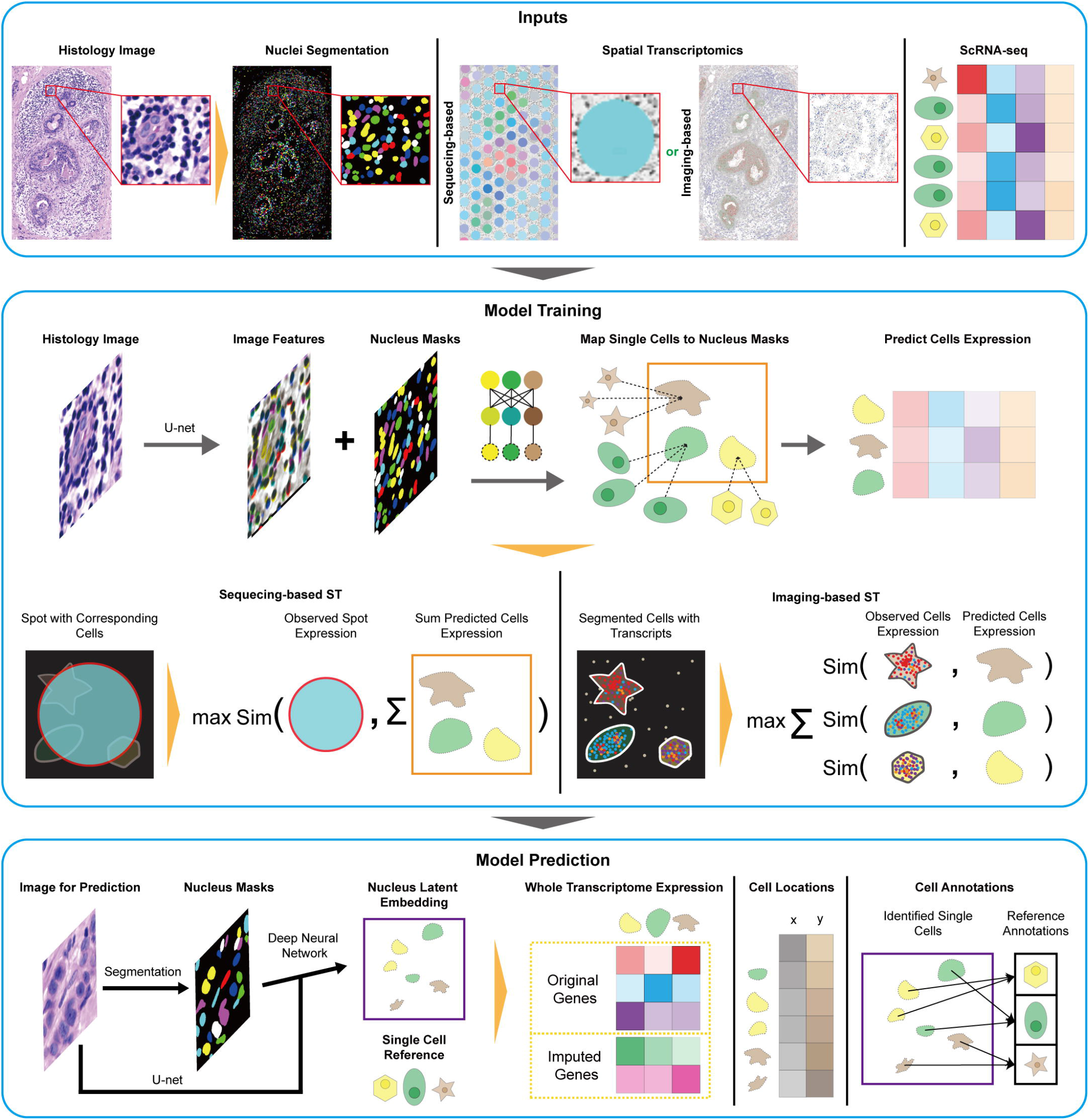
The Architecture of RedeHist. The overall architecture of RedeHist model encompasses model inputs, the processes of model training and prediction.

To train the RedeHist model, U-net is firstly applied to extract histology features. These features, along with nucleus masks, are fed into a neural network to produce latent embeddings for the masked regions. In addition, latent embeddings for single-cell references are also generated through fully connected layers. Based on these latent features, our model generates the cell abundance matrix employing a formula akin to scaled dot-product attention mechanism^8^. For imaging-based ST, the predicted expression matrix for mask regions is generated by performing a matrix multiplication of the cell abundance matrix and the single-cell expression matrix. For sequencing-based ST, RedeHist predicts the expression for masked regions, then sum the corresponding masked expressions for spots to generate predicted expression matrix for spots. Finally, our model maximizes cosine similarity between predicted and experimental observed expression matrix.

To predict transcriptomic profiles at single-cell resolution on histology images, image patches, along with their corresponding nucleus masks, are fed into the trained RedeHist model to generate a predicted expression matrix for single cells. Furthermore, RedeHist calculates the average spatial coordinate of nucleus masks for each cell to represent their corresponding spatial positions. Finally, the process of label transfer is undertaken to produce annotations of cell type for single cells.

### Implementation of RedeHist on Sequencing-based Spatial Transcriptomics

To assess the feasibility of training the RedeHist model on sequencing-based ST data, which capture whole transcriptomic expression at the spot level, we implemented our algorithm on a 10x Genomics Visium human breast cancer dataset. The Visium data, which captured 18,536 genes in 4,992 spots, was collected from 5 um tissue sections that had undergone H&E staining prior to sequencing. In addition, scFFPE-seq data was obtained from 2×25 um FFPE curls located adjacent to the tissue sections utilized for Visium^3^. The H&E image, Visium, and scFFPE-seq data were employed to predict ST at single-cell resolution, alongside cell annotation through RedeHist. The results illustrated distinct spatial distribution patterns among major cell types, while also revealing consensus patterns between the cell types and their corresponding marker genes (Fig. 2a). As we expected, ductal carcinoma in situ (DCIS) cells with marker gene *CEACAM6* were predominantly observed within ducts, whereas tumor cells expressing *EPCAM* breached the ductal architecture (Fig. 2a). T cells expressing *CD3E* gene often surrounded tumor or DCIS cells, while stromal cells expressing *POSTN* were widely distributed within the whole tissue (Fig. 2a). These results indicate RedeHist is a powerful tool for expression prediction and cell annotation based on histology images with sequencing-based ST data.

**Fig 2.**
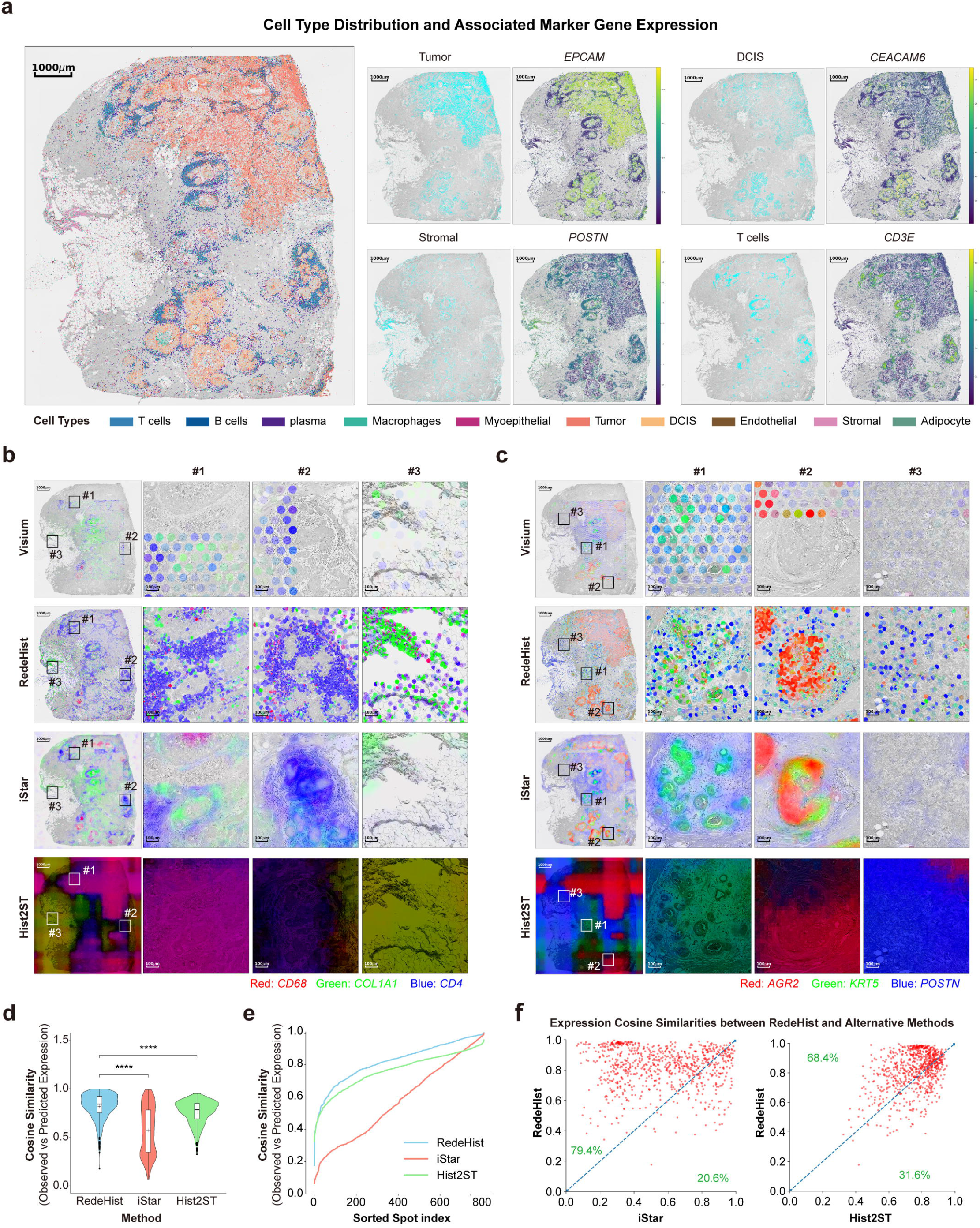
Implementation of RedeHist on Sequencing-based Spatial Transcriptomics. **a**. Cell type distribution from RedeHist results, illustrated using different colors (left). Spatial distribution of tumor, DCIS, stromal and T cells with their corresponding marker genes *EPCAM, CEACAM6, POSTN* and *CD3E* respectively (right). **b**. Distribution of *CD68, COL1A1*, and *CD4* from the Visium experiment, alongside prediction results of RedeHist, iStar, and Hist2ST. Illustrations at full scale or within three ROIs are provided from left to right. **c**. Distribution of *AGR2, KRT5*, and *POSTN* from the Visium experiment, alongside prediction results of RedeHist, iStar, and Hist2ST. Illustrations at full scale or within three ROIs are provided from left to right. **d**. Violin and box plot of cosine similarities between observed and predicted expression profiles for RedeHist and alternative approaches. The center line and the bounds of box refer to median, Q1 and Q3 of scores and the whisker equal to 1.5*(Q3-Q1). The minimum and maximum scores refer to Q1-whisker and Q3+whisker. The significance level marker denotes the level of significance under the null hypothesis. “*”, “**”, “***” and “****” denote significance levels of less than 0.05, 0.01, 0.001, and 0.0001, respectively. **e**. Line chart of cosine similarities between observed and predicted expression profiles. **f**. Scatter diagrams illustrate pairwise comparisons between RedeHist and other methods concerning spot-level expression similarities. Each point represents one spot.

To further demonstrate advantages of our approach, we compared the results obtained from RedeHist with those from the SOTA algorithms iStar^6^ and Hist2ST^5^, alongside the original Visium outputs. We firstly investigated the distributions of *CD68, COL1A1* and *CD4*, which serve as marker genes for macrophages, stromal cells, and T cells, respectively (Fig. 2b). We selected three regions of interest (ROIs) to demonstrate the superiority of RedeHist. In the ROIs #1 and #2, RedeHist accurately identified expression patterns for immune and stromal cells at single-cell resolution. IStar identified clear distribution patterns, but overlooked numerous immune cells in #1 and failed to achieve single-cell resolution in either #1 or #2. In the ROI #3, RedeHist successfully predicted gene expression in both high-density and low-density cell regions, spanning from the top to the bottom of the ROI, while iStar notably missed the predictions, particularly in the low-density cell regions. Furthermore, Hist2ST failed to identify expression patterns in these ROIs and the original Visium data only captured low-resolution gene expression patterns in specific regions due to the technical constraints in resolution and detection area (Fig. 2b). We then investigated the distributions of *AGR2, KRT5* and *POSTN*, the marker genes for tumor, myoepithelial and stromal cells respectively (Fig. 2c). Consistent with previous results, RedeHist accurately predicted expression at single-cell resolution in three ROIs. IStar exhibited limitations on both resolution (Fig. 2c #1 and #2) and prediction accuracy in low-density cell regions (Fig. 2c #3). Hist2ST and the original Visium data showed the same drawback as observed before.

Finally, we conducted a quantitative evaluation of all approaches by comparing expression similarity between the predictions derived from alternative methods and the observations obtained from the Visium experiment in testing dataset (See Methods). The results demonstrated that RedeHist yielded the most accurate predictions, attaining an average and median similarity of 0.8176 and 0.8408, respectively. In contrast, Hist2ST displayed an average and median performance of 0.7593 and 0.7844, while iStar recorded an average and median of 0.5697 and 0.5665 (Fig. 2d,e). Moreover, RedeHist outperformed iStar and Hist2ST on nearly 80% and 70% of spots respectively (Fig. 2f).

In conclusion, when trained on sequencing-based ST data, RedeHist not only achieves predictions at single-cell resolution but also surpasses SOTA approaches in terms of prediction accuracy.

### Implementation of RedeHist on Imaging-based Spatial Transcriptomics

In contrast to sequencing-based ST, imaging-based ST naturally achieves single-cell resolution but is often limited by the number of genes recovered. The design of RedeHist, incorporating nuclei segmentation results and scRNA-seq data, facilitates better utilization of single-cell resolution information and holds potential for recovering undetected genes. In this study, we applied RedeHist to a Xenium human breast cancer dataset that was adjacent to the tissue sections used for Visium and scFFPE-seq mentioned above. Thus, we not only predicted cell expression and performed cell annotation on H&E image similar to what we have done on the Visium dataset, but also imputed expression for undetected genes by employing scRNA-seq data. Our results showed the heterogeneous distribution of cell types, as well as the consistency between these cell types and their corresponding original and imputed marker genes (Fig. 3a). For example, myoepithelial cells expressing marker genes *KRT5* and *COL17A1* were frequently observed along the ducts, while endothelial cells with marker genes *VWF* and *CDH5* were found in close proximity to tumor and DCIS cells (Fig. 3a).

**Fig 3.**
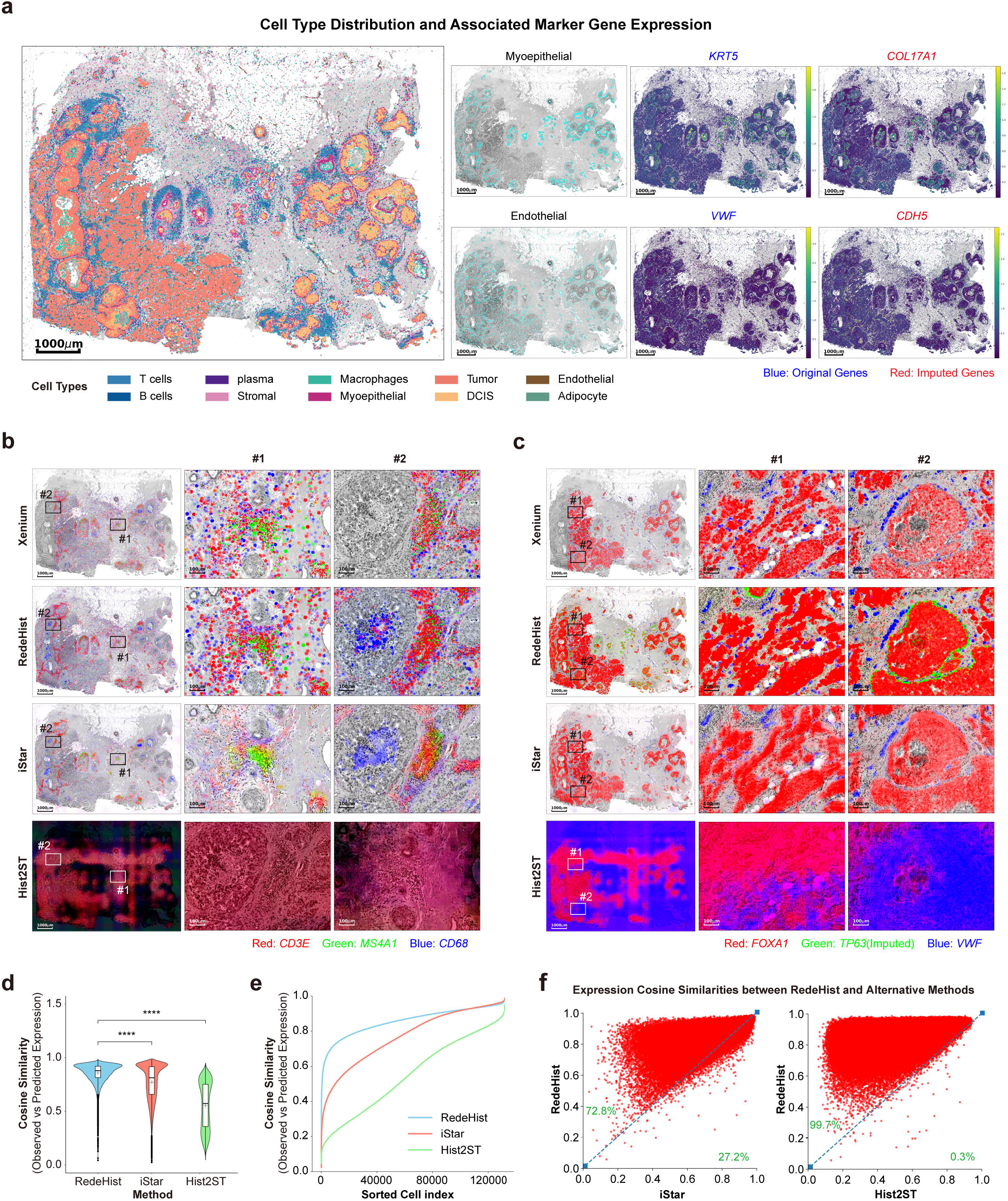
Implementation of RedeHist on Imaging-based Spatial Transcriptomics. **a**. Cell type distribution from RedeHist results, illustrated using different colors (left). Spatial distribution of myoepithelial and endothelial cells with original detected marker genes *KRT5* and *VWF* and imputed marker genes *COL17A1* and *CDH5* respectively (right). **b**. Distribution of *CD3E, MS4A1*, and *CD68* from the Xenium experiment, alongside prediction results of RedeHist, iStar, and Hist2ST. Illustrations at full scale or within two ROIs are provided from left to right. **c**. Distribution of original detected genes *FOXA1* and *VWF* and imputed gene *TP63* from the Xenium experiment, alongside prediction results of RedeHist, iStar, and Hist2ST. Illustrations at full scale or within two ROIs are provided from left to right. **d**. Violin and box plot of cosine similarities between observed and predicted expression profiles for RedeHist and alternative approaches. The center line and the bounds of box refer to median, Q1 and Q3 of scores and the whisker equal to 1.5*(Q3-Q1). The minimum and maximum scores refer to Q1-whisker and Q3+whisker. The significance level marker denotes the level of significance under the null hypothesis. “*”, “**”, “***” and “****” denote significance levels of less than 0.05, 0.01, 0.001, and 0.0001, respectively. **e**. Line chart of cosine similarities between observed and predicted expression profiles. **f**. Scatter diagrams illustrate pairwise comparisons between RedeHist and other methods concerning cell-level expression similarities. Each point represents one cell.

Next, we explored the feasibility of training SOTA models using imaging-based ST data and their applicability in predicting ST on histology images. To implement these approaches on Xenium dataset, we generated pseudo-spot and image patches for the model training (See Methods), followed by predictions on the entire images. We investigated the prediction results for *CD3E, MS4A1*, and *CD68*, which respectively represent T cells, B cells, and macrophages (Fig. 3b). In the Xenium experiment and RedeHist results, T cells and B cells were intermingled, whereas the iStar results revealed distinct aggregation areas for these cells (Fig. 3b #1 and #2). Similar to the findings in the Visium dataset, Hist2ST failed to identify expression patterns in all ROIs, and Xenium exhibited limitations in detection coverage. Furthermore, we investigated the expression of *FOXA1, TP63*, and *VWF* that representing tumor/DCIS cells, myoepithelial cells, and endothelial cells, respectively (Fig. 3c). The results from the Xenium experiment and RedeHist predictions revealed numerous independent endothelial cells adjacent to the invasive tumor, whereas iStar missed this observation, possibly due to low resolution (Fig. 3c #1). Furthermore, only RedeHist was able to impute the undetected gene *TP63* for myoepithelial cells and observed its expression along ducts (Fig. 3c #2).

For quantitative evaluation, we predicted the expression for each segmented cell using alternative approaches and compared it with the expression observed in the Xenium experiment (See Methods). RedeHist made the best prediction again with an average and median similarity of 0.8568 and 0.8778, respectively. IStar also performed well, achieving an average and median similarity of 0.7714 and 0.8125. However, the performance of Hist2ST has declined with an average and median similarity of 0.5521 and 0.5716 (Fig. 3d,e). Moreover, RedeHist outperformed iStar and Hist2ST on over 72% and 99% cells respectively (Fig. 3f). In conclusion, RedeHist not only fully utilized cell-level expression in imaging-based ST data to enhance prediction performance at single-cell resolution, but also enabled the imputation of undetected genes to obtain whole transcriptome expression profiles.

## Discussion

ST characterizes gene expression profiles while maintains spatial tissue context, thereby offering profound insights to elucidate the spatial organization and functional heterogeneity of gene expression within tissues and cells, which is pivotal for investigating diverse disease contexts and assessing value of clinical and translational application^19,20^. However, the clinical application of ST is constrained by high costs and labor-intensive procedures. Alternatively, predicting ST from histology images presents a cost-effective and readily accessible approach for researchers and clinicians, offering a valuable resource for diverse studies and facilitating the integration of histological data into transcriptional analysis. In addition, current ST technologies also suffer from limitations on resolution, number of genes recovered or detection range etc.

In this study, we develop RedeHist for ST prediction, cell annotation and gene imputation at single-cell resolution based on histology images. On one hand, we employ deep neural network for feature extraction and nuclei segmentation algorithms for cell identification to improve the predictive resolution. On the other hand, inspired by deconvolution algorithms, we also employ scRNA-seq data and integrate them with predicted cell expression profiles, which enables our algorithm to impute undetected genes and improve prediction accuracy. The applications of RedeHist to both sequencing-based and imaging-based ST datasets demonstrates its applicability for ST prediction, cell annotation, and gene imputation from histology images, outperforming SOTA algorithms in terms of predictive accuracy and resolution.

Although RedeHist offers significant advantages, further work remains to be undertaken. Firstly, owing to the high resolution and whole transcriptome profiles coupled with spatial information provided by RedeHist’s predictions, there is a compelling interest in exploring biological discoveries hidden in the data, but the data complexity raises an unmet analytical challenge. Secondly, additional research is needed to test and demonstrate the potential utility of spatial transcriptomics prediction for large-scale, multicenter, cross-organ histology images. In summary, we have developed a novel spatial transcriptomics prediction tool RedeHist and demonstrated the functionality and performance. RedeHist may serve as a valuable tool for bridging the application of histology images and ST technologies across diverse research disciplines.

## Acknowledgements

This work was supported by Changping Laboratory, the National Natural Science Foundation of China (32022016 X.R., 92159305 X.R., and 31991171X.R.), National Key R&D Program of China (2020YFE0202200 X.R. and 2022YFC3400904 X.R.).

## Author Contributions Statement

X.R. conceived this study, designed the algorithm, supervised the analysis, and wrote the manuscript. Y.Z. developed the algorithm and the software, conducted the data analysis, and wrote the manuscript. J.Z conducted the data analysis and wrote the manuscript.

## Methods

### Model Overview

RedeHist employs deep neural network for predicting single cell expression on histology images. In detail, the histology images underwent an initial pre-processing stage utilizing an automatic nuclei segmentation algorithm for identifying nucleus masks. RedeHist then incorporates the original images, along with the nucleus masks and scRNA-seq references, to train a robust deep neural network model. The trained model can subsequently be applied for predicting expression at single-cell resolution in any histology images.

### Pre-processing

Pre-processing module is designed to produce input data available for RedeHist model. The original histology image can be represented as a three-dimensional tensor (width, height, channel). We first generate a nucleus masks matrix by inputting the tensor into a nuclei segmentation algorithm. This matrix uniquely represents the identity of each nucleus.

After segmentation, RedeHist divides the histology image tensor into patches of size d*d. For sequencing-based ST, the quantity of image patches is equal to the number of spots, with the patch centers aligning precisely with the spot centers. Moreover, we set one image patch (spot) corresponding to m nucleus masks. In each iteration, RedeHist randomly selects m nuclei within the spot without repetition and generates m nucleus patches by capturing the same region from the corresponding image patch. For each nucleus patch, it assigns a value of one to corresponding nucleus mask pixels and zero to other pixels. If the number of nuclei within a spot is less than m, the remaining nucleus patches are set to zero. For imagining-based ST, the number of image patches equals the count of nuclei within the ST capture region, with patch centers corresponding to nucleus centers. Nucleus patches are generated following the same process as in sequencing-based ST, with the exception of setting m to 1. In addition, we extend the boundaries of nuclei outward until either a maximum distance of 15um is reached or the boundary of another cell is encountered, thereby encompassing cell regions. We then count the transcripts within these regions to construct the expression matrix for cells.

We finally obtain (1) image patch tensor X of dimensions n×d×d×c, n and c represent the number of patches and channels respectively; (2) nucleus masks tensor Y of dimensions n×d×d×m; (3) expression matrix E for ST of dimensions n×g, where g is the number of intersection genes between ST and scRNA-seq; (4) expression matrix S for scRNA-seq of dimensions k×g and the genes in E and S are in same order. For training process, we implement a batch training strategy where the first dimension of input tensors X, Y, and E is b for each iteration, with b representing the batch size.

### Model Structure

Using the image patch tensor X as input, RedeHist applies U-net to extract pixel-level histological features, followed by a linear layer with layer normalization for feature transformations. The output dimension for both the U-net and linear layer for each pixel is f.

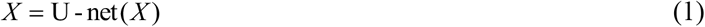

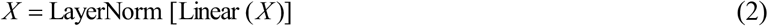

We then employs pixel-level features with nucleus masks tensor Y to generate patch-level features. In order to match the dimensions, X and Y are replicated m and f times respectively, where the dimensions become b×d×d×m×f. Thus, patch feature tensor Z of dimensions b×m×f was generated.

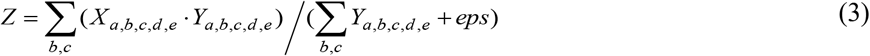

We also integrate an attention mechanism to enhance the patch features. Tensor Z is replicated d^2^ time to become dimension of b×d×d×m×f. Then, pixel-level attention weight tensor W was generated.

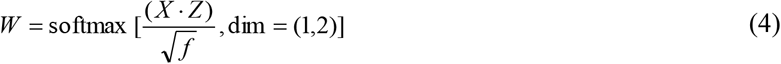

Based on the attention weight, the patch feature tensor Z is updated.

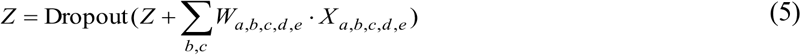

For scRNA-seq data, we apply 3 liner layers with active function (6)∼(8) to encode scRNA-seq expression matrix S and the dimension of input and output features for these layers are 200 except the input dimension for the first layer is g.

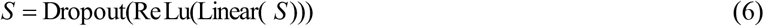

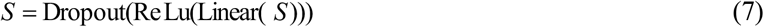

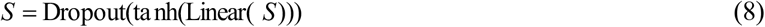

We then predict cell abundance for each nuclei nucleus mask region. The idea is similar to deconvolution ST data using scRNA-seq reference. We suppose that the expression of each cell in histology images can be represented as the sum of multiple single-cell expressions with varying abundances. Hence, we predict cell abundances for nuclei on histology images through (9) and obtain cell abundance matrix A of dimension b×m×k. The matrix A undergoes an additional normalization procedure to guarantee that the total sum of cell abundances within each nucleus region equal to 1.

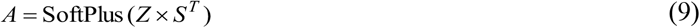

Due to batch effect and dropout in both ST and scRNA-seq data, we generated adjust scRNA-seq matrix S’ to bridge the gap between ST and scRNA-seq data.

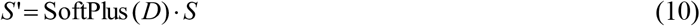

Where, the matrix D is a trainable matrix initialized with unity values and shares the same dimensions as matrix S. Thus, the expression matrix P of dimension b×m×g for nucleus masks on batch images could be predicted by multiplying the cell abundance matrix A with the adjust scRNA-seq expression matrix S’.

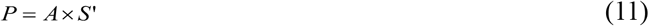

As in each spatial spot, the observed expression denotes the cumulative expression derived from multiple single cells. Therefore, we aggregate the predicted expression values from m nucleus masks within the corresponding patch image, resulting in the generation of a spot expression prediction matrix E’ of dimensions b×g.

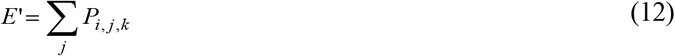

The expression similarity between observed and predicted for spots could be calculated and the corresponding first loss function is give in (13).

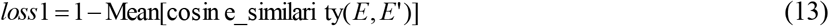

Additionally, the secondary loss function is optimized to enhance the similarity between the original and adjusted scRNA-seq data.

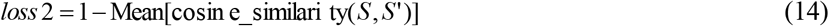

We also minimize two entropy losses. The first one ensures that the predicted abundance for each nucleus more likely originates from one cell type, while the second promotes a more uniform distribution of the overall abundance across different cell types. According to cell type annotation of scRNA-seq data, we generate cell type proportion matrix B of dimension b×m×c according to A, where c refer to number of cell type. Then, we compute the two entropy as:

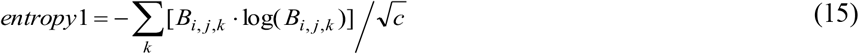

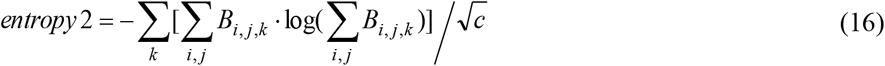

The dimensionality of entropy1 is b×m, whereas entropy2 is a scalar. With reference to these entropy terms, the third and fourth losses are presented in equations (17) and (18) respectively.

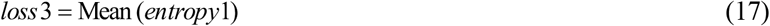

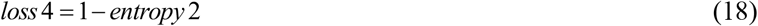

Eventually, the overall loss function is given as:

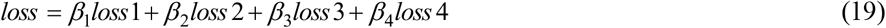

The β values in equation (19) represent the weights assigned to each individual loss function.

### Model Training

Given imaging-based or sequencing-based ST data accompanied by high-resolution histology images and scRNA-seq references, we preprocess the data in accordance with the procedures delineated in the “Pre-processing” section. Specifically, we set m to 1 for imaging-based ST and 20 for sequencing-based ST, respectively. To minimize the loss function described in equation (19), we utilize Adam as the optimizer and specify a learning rate of 0.001 for D in equation (10), while setting the learning rate to 0.00005 for the remaining parameters. To conduct batch training, we specify the number of epochs as 50 and batch size as 16 or 4 for imaging-based or sequencing-based ST respectively. We also set dropout ratio to 0.3 to avoid over-fitting.

### Model Prediction

After completion of model training, the model is capable of predicting the expression of each cell within histology images. Upon receiving a novel image, the initial step involves cell segmentation to acquire nucleus mask regions. Subsequently, the model maps the scRNA-seq reference to these regions, generating an abundance matrix for the designated regions. Eventually, the model predicts expression for each region, where the predicted expression serves as a representation of the expression of the corresponding cell in the histology images. The dropout ratio used in the prediction process is 0.

### Generation of ScRNA-seq Reference of Human Breast Cancer

We downloaded scFFPE-seq data from 10X Genomics website, yielding a total number of 30,365 single-cell profiles. Due to absence of adipocyte, we incorporated an additional snRNA-seq dataset containing 25,871 adipocyte single-cell profiles sourced from the Single-Cell Atlas of Human Adipose Tissue^21^. Consequently, our combined single cell reference comprises a comprehensive total number of 56,236 single-cell transcriptomes.

### Pseudo-spot for Xenium Dataset

It is worth noting that comparable approaches (e.g. iStar and Hist2ST) often overlook nuclei segmentation and fail to incorporate nucleus masks during the training process. So, they cannot be directly applied for training imaging-based ST data (e.g. Xenium and MERFISH). In this study, we acquired the Xenium dataset from the 10X Genomics website and generated pseudo-spot for Xenium data to facilitate training of comparable models. We divided the H&E image that overlaps with the Xenium capture area into 128×128 patches and subsequently tallied all transcripts present within each patch to produce an expression profile. The pseudo dataset we generated comprises images and center coordinates of patches, along with their corresponding expression profiles, which were employed for training alternative models.

### Application of RedeHist to different ST Datasets

We implemented RedeHist on an imaging-based Xenium dataset and a sequencing-based Visium dataset, both pertaining to human breast cancer specimens. The RedeHist model was trained using the comprehensive reference of scRNA-seq data that we generated, along with H&E images and spatial transcriptomics (ST) data as input. For model parameters, we set d values of patch sizes to 192 and 128 for Visium and Xenium respectively, while maintaining the default settings for all other parameters. To determine the gene set for training, we calculated the overlap of genes between the Xenium transcripts or Visium data and the scRNA-seq reference. In the case of Xenium ST data, all 313 overlapping genes were utilized for model training. Regarding the Visium ST data, 2,000 highly variable genes which were selected by ROGUE^22^ on the scRNA-seq reference were employed for the same purpose. After training the model, we predicted expression profiles for single cells on H&E images and identified the corresponding cell types by taking into account the most prevalent cell type within the cell abundance matrix.

### Application of iStar to different ST Datasets

IStar was implemented on both Visium and Xenium datasets. For the Visium dataset, the model was trained on a dataset identical to RedeHist, encompassing consensus highly variable genes, H&E images, and ST data. Recognizing the divergence between the iStar demo dataset and our own, we adjusted the pixel size to 0.37 and the radius to 64 to align them precisely with our image specifications. For the Xenium dataset, we trained the model utilizing the aforementioned pseudo-spot, which comprised 313 genes. Specifically, we adjusted the pixel size to 0.4 and the radius to 64, while maintaining all other parameters as default. We finally obtained pixel-level prediction results for individual genes and subsequently elevated the resolution to ensure spatial scale consistency.

### Application of Hist2ST to different ST Datasets

Hist2ST was implemented on both Visium and Xenium datasets using 128×128 patches paired with corresponding expression profiles and the same gene set as in RedeHist for consistency. During the training process, we encountered a “Not a Number (NaN)” error while using the original spatial coordinates, both with and without the application of the StandardScaler from sklearn. So, we finally scaled the spatial coordinates by dividing them by 1,000. Moreover, we set fig_size equal to 128, dropout ratio equal to 0.2, while maintaining the default values for the remaining parameters. After training the model, the whole H&E image was divided into 128×128 patches with a step size of 64 pixels. Hist2ST predicted the expression profiles for each patch and we then merged the results and eventually generated expression map for the whole image.

### Benchmarking

To quantitatively evaluate performance of all approaches, we compared expression similarity between ground truth and the approaches on both the Visium and Xenium datasets. For the Visium dataset, we trained the model using expression profiles from a randomly selected set of 3,992 spots (training dataset) for all three methods. Subsequently, we employed the model to predict spot-level expression profiles in the remaining 1,000 spots (testing dataset). For the Xenium dataset, we firstly employed expanding approaches to generate cell regions and cell masks from nuclei segmentation results^3^. Then, cell-level prediction results were generated for alternative models by summing pixel-level prediction results within cell masks. Eventually, we calculated spot-level and cell-level cosine similarity for the Visium and Xenium datasets, respectively, between predicted and observed expression profiles. A higher cosine similarity indicates that the prediction results are more realistic and accurate.

## Data Availability

Both scRNA-seq, Visium, and Xenium data, as well as H&E images of human breast cancer were available at https://www.10xgenomics.com/products/xenium-in-situ/preview-dataset-human-breast. SnRNA-seq data of human adipose tissue was available on the NCBI GEO database with accession number GSE176171.

## Code Availability

The package is available on github with detailed documentation (https://github.com/Roshan1992/RedeHist).

